# Symbiotic nutrient cycling enables the long-term survival of Aiptasia in the absence of heterotrophic food sources

**DOI:** 10.1101/2022.12.07.519152

**Authors:** Nils Rädecker, Anders Meibom

## Abstract

Phototrophic Cnidaria are mixotrophic organisms that can complement their heterotrophic diet with nutrients assimilated by their algal endosymbionts. Metabolic models suggest that the translocation of photosynthates and their derivatives from the algae may be sufficient to cover the metabolic energy demands of the host. However, the importance of heterotrophy to the nutritional budget of these holobionts remains unclear. Here, we report on the long-term survival of the photosymbiotic anemone Aiptasia in the absence of heterotrophic food sources. Following one year of heterotrophic starvation, these anemones remained fully viable but showed an 85 % reduction in biomass compared to their regularly fed counterparts. This shrinking was accompanied by a reduction in host protein content and algal density, indicative of severe nitrogen limitation. Nonetheless, isotopic labeling experiments combined with NanoSIMS imaging revealed that the contribution of algal-derived nutrients to the host metabolism remained unaffected due to an increase in algal photosynthesis and more efficient carbon translocation. Taken together, our results suggest that, on a one-year timescale, heterotrophic feeding is not essential to fulfilling the energy requirements of the holobiont. But, while symbiotic nutrient cycling effectively retains carbon in the holobiont over long time scales, our data suggest that heterotrophic feeding is a critical source of nitrogen required for holobiont growth under oligotrophic conditions.

## Introduction

Photosymbiotic Cnidaria, such as corals and anemones, dominate shallow hard-bottom substrates in the oligotrophic tropical ocean (Pandolfi 2002). The key to their evolutionary and ecological success under these conditions lies in their association with endosymbiotic algae of the family Symbiodiniaceae (Stanley 2006; Stanley and van de Schootbrugge 2009). Efficient nutrient exchange in these symbioses couples the heterotrophic metabolism of the host with the autotrophic metabolism of their algal symbionts (Yellowlees et al. 2008; Cunning et al. 2017). Consequently, photosymbiotic Cnidaria are considered mixotrophic as they can acquire nutrients via heterotrophy and autotrophy alike (Fox et al. 2018; Radice et al. 2019). Under oligotrophic conditions, this confers an ecological advantage that enables these animals to outcompete other benthic organisms restricted to either heterotrophic or autotrophic nutrient sources (Muscatine and Porter 1977; McCook 2001).

In the stable symbiosis, the algal symbionts translocate a large proportion of their photosynthates in the form of sugars and sterols to their host (Falkowski et al. 1984; Burriesci et al. 2012; Tremblay et al. 2014; Hambleton et al. 2019). This carbon translocation fuels the host metabolism and may be sufficient to cover the host’s energy demand under optimal environmental conditions (Davies 1984; Rinkevich 1989; Tremblay et al. 2012). The translocated photosynthates have been referred to as ‘junk food’ because their low nitrogen content limits their potential for anabolic incorporation (Falkowski et al. 1984; Dubinsky and Jokiel 1994). Hence, the utilization of photosynthates by both symbiotic partners depends, in part, on their access to inorganic nitrogen sources from the surrounding seawater (Davies 1984; Morris et al. 2019; Rädecker et al. 2021). However, under the oligotrophic conditions that prevail in the tropical ocean inorganic nitrogen availability is limited (O’Neil and Capone 2008).

In contrast, heterotrophic nutrient sources have a proportionally higher nitrogen content that enables efficient anabolic assimilation (Hughes et al. 2010). There is ample evidence demonstrating the nutritional benefits of heterotrophic feeding, e.g., in the form of organic nitrogen or vitamins for both symbiotic partners (Goreau et al. 1971; Porter 1976; Houlbrèque and Ferrier-Pagès 2009). For example, high rates of heterotrophic feeding seem to enable corals to compensate for reduced algal-derived nutrient availability during bleaching; i.e., the stress-induced breakdown of the cnidarian-algal symbiosis (Grottoli et al. 2006; Anthony et al. 2009). However, as our understanding of potential prey dynamics (e.g., zooplankton abundance) and cnidarian grazing on coral reefs remains limited (Lowe and Falter 2015), the importance of heterotrophic nutrients for sustaining the stable cnidarian-algal symbiosis is less clear.

Here, we performed a starvation experiment using the photosymbiotic sea anemone Aiptasia to study the role of heterotrophic nutrient acquisition in symbiosis. For this, we reared Aiptasia for one year in the absence of any heterotrophic nutrient sources. This permitted us to examine the effects of heterotrophic starvation on the symbiosis in light of the underlying carbon and nitrogen cycling and explore the limits of autotrophic nutrient acquisition in the photosymbiotic Cnidaria.

## Material & Methods

### Animal husbandry & experimental design

The experiments and measurements were performed on the photosymbiotic cnidarian model organism Aiptasia, i.e., *Exaiptasia diaphana* (Puntin et al. 2022). We used the clonal host line CC7 with its native algal symbiont community dominated by the *Symbiodinium linucheae* strain SSA01 (Sunagawa et al. 2009; Grawunder et al. 2015). Animal cultures were reared in 2 L acrylic food containers filled with artificial seawater (35 PSU, Pro-Reef, Tropic Marin, Switzerland). Artificial seawater was freshly prepared in the dark to minimize any potential microbial contamination. Culture stocks were kept at a constant temperature of 20°C under a 12 h: 12 h light-dark cycle (photosynthetic active radiation = 50 μE m^-2^ s^-1^) using an Algaetron 230 incubator (Photo System Instruments, Czech Republic). Animals were fed once a week with freshly hatched *Artemia salina* nauplii (Sanders GSLA, USA) followed by a complete water exchange and removal of biofilms.

For the experiment, all animals were reared under the same conditions as outlined above for one year. However, while half of the animals (two culture containers with five animals each) were fed weekly with *Artemia* nauplii (regularly fed control), the other half (two culture containers with five animals each) was reared in the absence of any food sources (heterotrophically starved). Apart from this, all culturing parameters were kept identical, including the weekly cleaning and water exchange. Pedal lacerates of Aiptasia were continuously removed to avoid experimental biases due to differences in asexual reproduction and resulting densities of animal populations.

### Phenotypic characterization

Following the one-year experiment, treatment responses were recorded. First, photos of representative phenotypes for each of the treatments were taken with an OM-1 camera and a 60 mm f2.8 macro-objective (OM System, Japan) using identical illumination and exposure settings. Then, three animals were collected from each treatment group, transferred to a pre-weighed 1.5 mL Eppendorf tube, and homogenized in 500 μL Milli-Q water using a PT1200E immersion dispenser (Kinematica, Switzerland).

Host and algal symbiont fractions were immediately separated by centrifugation (3000 g, 3 min, sufficient to remove > 95 % of algal symbionts from the supernatant) and the host supernatant was transferred into a new pre-weighed 1.5 mL tube, flash-frozen in liquid nitrogen, and stored at -20°C for later analysis. The algal symbiont pellet was resuspended in 500 μL Milli-Q water and rinsed by one additional centrifugation and resuspension step. Algal symbiont concentrations were quantified in three technical replicates per sample based on cell shape and chlorophyll autofluorescence using a CellDrop cell counter (DeNovix, USA). The protein content in the defrosted host supernatant was quantified in three technical replicates using the Pierce Rapid Gold BCA Protein Assay Kit (Thermo Scientific, USA) according to the manufacturer’s instructions. Algal concentrations and host protein content were extrapolated to the initial sample volume and normalized to holobiont biomass. Holobiont biomass was approximated as dry weight. For this, host and symbiont fractions were dried at 45°C until the weight was stable and the initial weight of empty tubes was subtracted from the final weight. The weight of host and symbiont fractions was corrected for aliquots taken for sample measurements to approximate the dry weight of the holobiont as a whole, i.e., host + symbiont fraction.

### Isotope labeling & sample processing

To study treatment effects on symbiotic interactions, we quantified inorganic carbon and nitrogen assimilation and translocation in the symbiosis. For this, three animals from each treatment were transferred to 50 mL glass vials. For isotopic labeling, vials were filled with minimal artificial seawater medium (35 PSU, pH 8.1, 355.6 mM NaCl, 46.2 mM MgCl_2_, 10.8 mM Na_2_SO_4_, 9.0 mM CaCl_2_, 7.9 mM, KCl; (Harrison et al. 1980)) containing 2.5 mM NaH^13^CO_3_ and 10 μM ^15^NH_4_Cl. In addition, one additional animal per treatment was transferred into a vial filled with minimal artificial seawater medium without heavy isotope tracers to serve as unlabeled controls for NanoSIMS measurements. Animals were incubated for 6 h in the light at their regular culture conditions before being transferred to a fixative solution (2.5 % glutaraldehyde and 1 % paraformaldehyde in 0.1 M Sorensen’s phosphate buffer). Samples were fixed for 1 h at room temperature followed by 24 h at 4°C before being stored in a preservative solution (1 % paraformaldehyde in 0.1 M Sorensen’s phosphate buffer) at 4 °C until further processing. Within four days of fixation, samples were dissected and individual tentacles were processed for resin embedding. Following secondary fixation in 1 % OsO_4_ for 1 h, samples were rinsed (3 x MiliQ for 10 min) and dehydrated in a series of increasing ethanol concentrations (30 % for 10 min, 50 % for 10 min, 2 × 70 % for 10 min, 3 × 90 % for 10 min, and 3 × 100 % for 10 min) before being transferred to acetone (100 % for 10 min). Dehydrated samples were gradually infiltrated with SPURR resin (Electron Microscopy Sciences, USA) at increasing concentrations (25 % for 30 min, 50 % for 30 min, 75 % for 1 h, and 100 % overnight) and the resin was polymerized at 65°C for 48 h. Embedded samples were cut into semi-thin sections (200 nm) using an Ultracut E ultramicrotome (Leica Microsystems, Germany) and transferred onto glow-discharged silicon wafers.

### Electron microscopy and NanoSIMS imaging

For scanning electron microscopy (SEM), sections were stained with uranyl acetate (1% for 10 min), followed by Reynolds lead citrate solution (10 min). Images of tentacle of sections were taken on a GeminiSEM 500 field emission scanning electron microscope (Zeiss) at 3 kV with an aperture size of 30 μm, and a working distance of 2.8 mm, using the energy selective backscatter detector (EsB; ZEISS) with the filter-grid set at 121 V. For NanoSIMS imaging, sections on wafers were sputter-coated with a 12 nm gold layer using an EM SCD050 (Leica Microsystems) and the samples were analyzed with a NanoSIMS 50L instrument (Hoppe et al. 2013). To remove the metal coating, target sample areas were pre-sputtered for 5 min with a primary beam of ca. 6 pA. Data were collected by rastering a 16 keV primary ion beam of ca. 2 pA Cs^+^ focused to a spot size of about 150 nm across the sample surface of 40 × 40 μm with a resolution of 256 × 256 pixels and a pixel dwell time of 5 ms; except for correlative SEM + NanoSIMS images that were recorded at 55 × 55 μm but not included in the data analysis. The secondary ions ^12^C_2-_, ^12^C^13^C^-, 12^C^14^N^-^, and ^12^C^15^N^-^ were simultaneously collected in electron multipliers at a mass resolution of about 9000 (Cameca definition), sufficient to resolve potentially problematic mass interferences. For each sample, seven to eight areas were analyzed in five consecutive image layers. The resulting isotope maps were processed using the ImageJ plug-in OpenMIMS (https://github.com/BWHCNI/OpenMIMS/wiki). Mass images were drift- and dead-time corrected, the individual planes were added and ^13^C/^12^C and ^15^N/^14^N maps were expressed as hue-saturation-intensity images, where the color scale represents the isotope ratio. ^13^C and ^15^N assimilation was quantified by drawing regions of interest (ROIs) of individual algal symbionts and surrounding host gastrodermis based on ^12^C^14^N^−^ maps. For this, algal symbiont ROIs were drawn by outlining individual algal cells and host ROIs were drawn in a circle with a 15 μm diameter around the centroid of the algal symbiont whilst excluding any algal symbionts and symbiosome content from the ROI. For unlabeled control Aiptasia, 80 host and 80 algal symbiont ROIs were analyzed across two animal replicates. For isotopically labeled Aiptasia, 120 host and 120 algal symbiont ROIs were analyzed across three animal replicates per treatment. Based on this, the ^13^C and ^15^N assimilation in individual ROIs was expressed as atom % excess (in comparison to corresponding ROIs of unlabeled control Aiptasia). Due to the clonal nature of Aiptasia and the identical environmental conditions of animals within the same treatment, individual ROIs were considered independent measurements across animal replicates for the purpose of this study.

### Statistical analyses

Treatment effects on phenotypic responses, i.e., biomass, host protein content, and symbiont density, were analyzed using two-sided unpaired Student’s *t*-tests. Isotope ratios from NanoSIMS analysis were log-transformed to meet model assumptions and analyzed with linear models (LM) using the respective symbiotic partner (host/symbiont) and treatment (fed/starved) as explanatory variables. To test individual differences between groups LMs were followed up with a Tukey HSD post hoc comparison.

## Results

### Holobiont biomass loss in the absence of heterotrophic nutrients

After one year of husbandry in the absence of heterotrophic food sources, no mortality was observed and Aiptasia remained viable but had ceased any detectable asexual propagation via pedal lacerates. Starved animals showed pronounced phenotypic differences compared to their regularly fed counterparts. Specifically, starvation resulted in a reduction in body size, a paler appearance, and a loss of 85 % of their dry weight (Fig. 1A,B; Student’s *t*-test, *t* = 4.71, *p* = 0.042). This decline in holobiont biomass was, at least in part, driven by a strong decline in host protein content and algal symbiont density, which both decreased by more than 80 % on average when normalized to holobiont biomass (Fig. 1C,D; for host protein: Student’s *t*-test, *t* = 5.39, *p* = 0.014; for algal symbiont density: Student’s *t*-test, *t* = 3.85 *p* = 0.047 for algal symbiont densities).

**Fig. 1.**
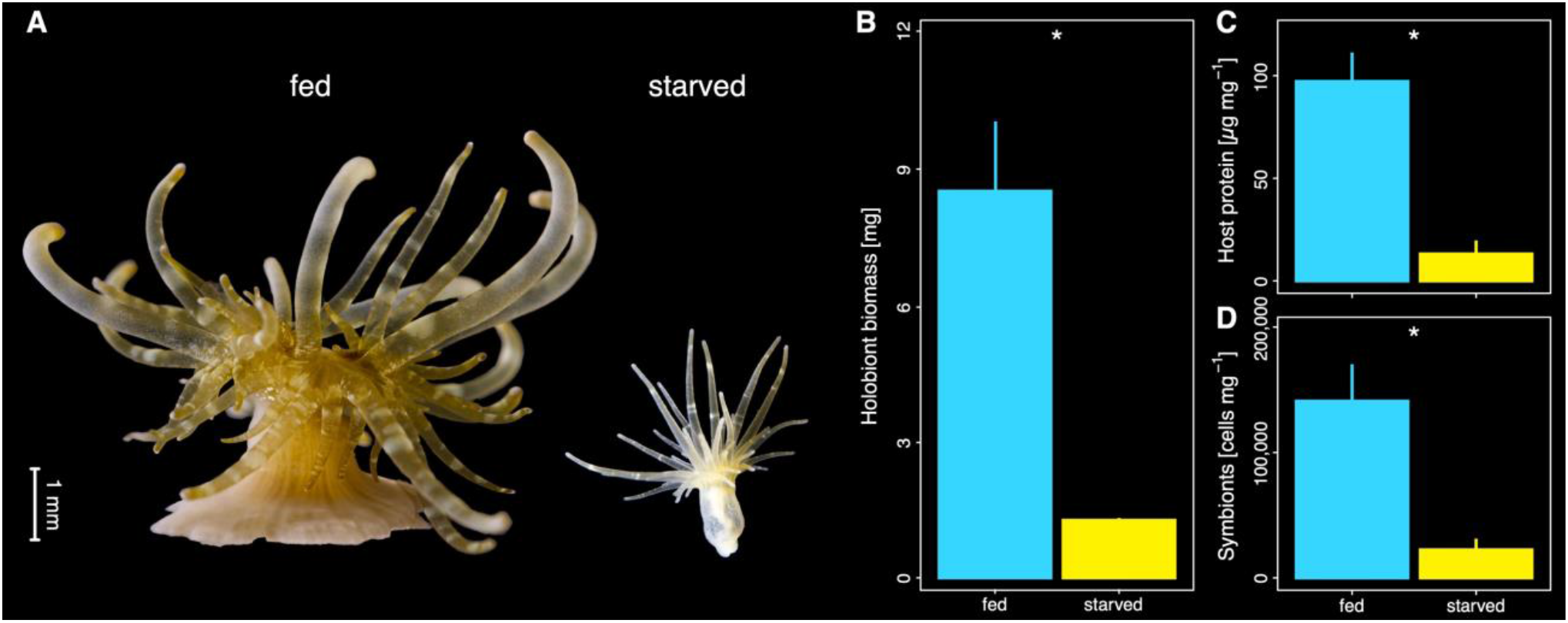
Phenotypic response to heterotrophic starvation in Aiptasia. **(A)** Representative photos illustrating the phenotype of animals that were regularly fed (left) or reared for one year without heterotrophic food sources (right). **(B)** Holobiont biomass expressed as dry weight of fed and starved Aiptasia. **(C)** Protein content of the host tissue per holobiont biomass of fed and starved Aiptasia. **(D)** Algal symbiont density per holobiont biomass of fed and starved Aiptasia. Three Aiptasia replicates were analyzed for each treatment. Asterisks indicate significant effects between treatments (*p < 0.05).

### Enhanced photosynthetic performance of algal symbionts sustains host metabolism during heterotrophic starvation

SEM images of tentacle sections confirmed that Aiptasia from both treatments hosted algal symbionts in their gastrodermal tissue. However, gastrodermal cells from starved Aiptasia appeared to contain a higher density of lipid bodies than those of their regular fed counterparts (Fig. 2A,B, Fig. S1). Likewise, algal symbionts from starved Aiptasia appeared to have a higher lipid content in their cells than those from regularly fed Aiptasia, albeit this trend was less clear than in the host tissue (Fig. S1).

**Fig. 2.**
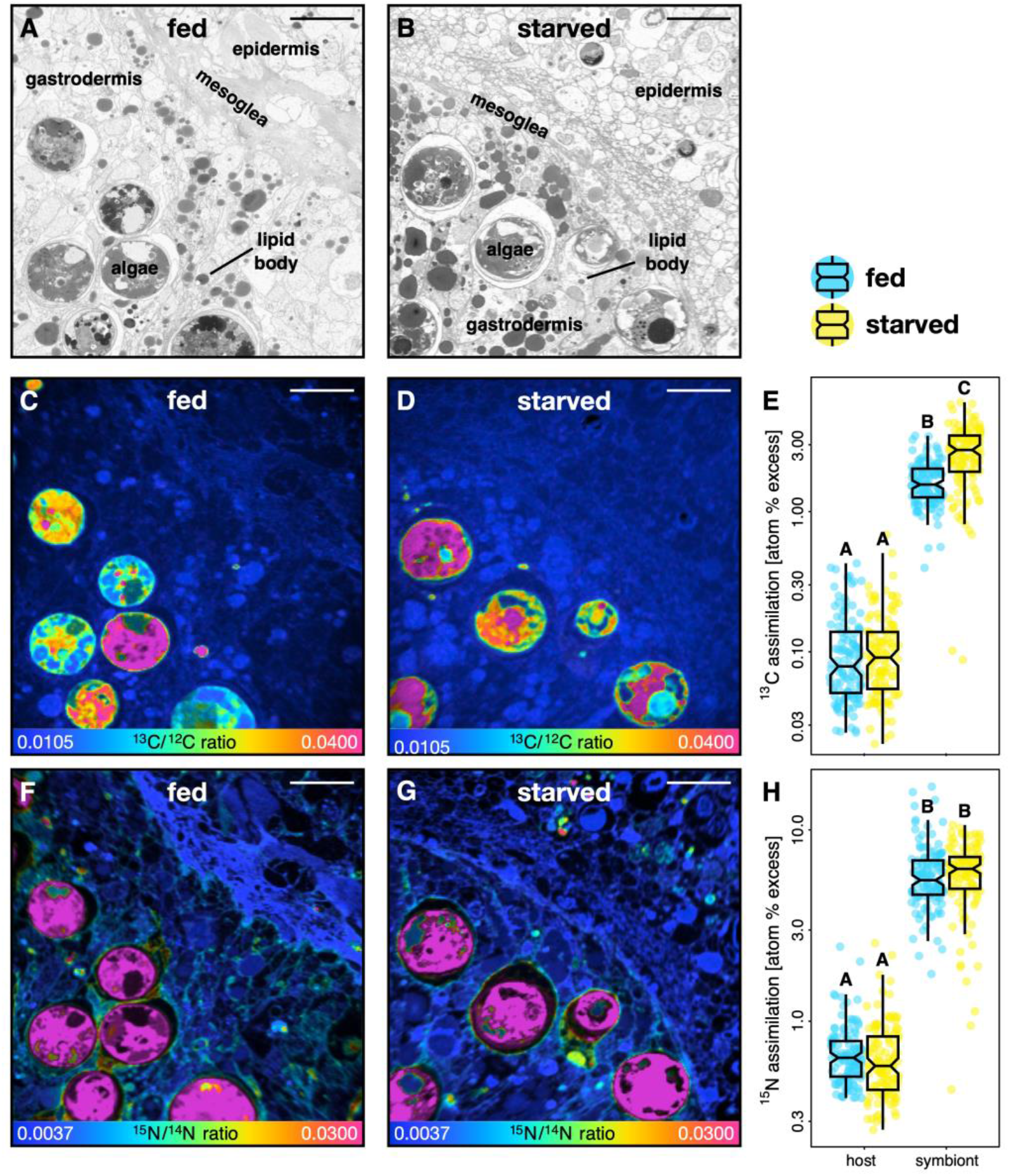
Correlative SEM and NanoSIMS imaging of symbiotic carbon and nitrogen cycling in fed and starved Aiptasia. **(A,B)** Representative SEM images of tentacle semi-thin sections of regularly fed and starved Aiptasia, respectively. **(C**,**D)** Corresponding NanoSIMS images illustrating H^13^CO_3-_ assimilation and translocation as ^13^C/^12^C isotope ratio maps in regularly fed and starved Aiptasia. **(E)** Boxplots and data points of ^13^C enrichment for the host gastrodermis and the algal symbiont cells. **(F**,**G)** Corresponding NanoSIMS images illustrating ^15^NH_4_^+^ assimilation as ^15^N/^14^N isotope ratio maps in regularly fed and starved Aiptasia. **(H)** Boxplots of ^15^N enrichment for the host gastrodermis and the algal symbiont cells. NanoSIMS ratio maps are shown as hue saturation images with blue representing no (or low) enrichment and pink representing the highest level of enrichment. Note the logarithmic scale for **E**,**H**. 120 regions of interest across three Aiptasia replicates were analyzed per symbiotic partner and treatment combination. Scale bars are 10 μm. Different letters above boxplots indicate significant differences between groups (p < 0.05).

NanoSIMS imaging revealed that metabolic interactions in the cnidarian-algal symbiosis remained remarkably stable during heterotrophic starvation despite the pronounced phenotypic response of the Aiptasia holobiont. Consistent with previous reports (Christophe Kopp et al. 2015; Rädecker et al. 2018), the ^13^C enrichment from ^13^C-bicarbonate assimilation/translocation was highest in the algal symbionts with host ^13^C enrichment primarily observed in localized hotspots corresponding to lipid bodies in the correlative SEM images (Fig. 2C,D, Fig. S2; host/symbiont differences: LM, F = 3189.76, *p* < 0.001). Despite drastic declines in algal symbiont densities during heterotrophic starvation, overall ^13^C enrichment remained stable in the gastrodermal tissue of the host (Tukey’s HSD, *p* = 0.951). This was best explained by the enhanced photosynthetic performance of algal symbionts, which exhibited an increase of nearly 50 % in ^13^C enrichment in starved animals (Fig. 2E, Fig. S2; Tukey’s HSD, *p* < 0.001).

The constant availability of photosynthates in the symbiosis during heterotrophic starvation was reflected in a maintained anabolic assimilation of ^15^N-ammonium by both symbiotic partners. Consistent with previous studies, the algal symbionts acquired the highest ^15^N enrichments from ammonium assimilation (Pernice et al. 2012; C. Kopp et al. 2013; Rädecker et al. 2018), but the host also exhibited clearly measurable ^15^N enrichments in both epidermal and gastrodermal tissue layers (Fig. 2F,G, Fig. S2; host/symbiont differences: LM, F = 3320.37, p < 0.001). Thus, heterotrophic starvation did not alter the ability for ammonium assimilation of either symbiotic partner (Fig. 2H, Fig. S2; Tukey’s HSD, *p* = 0.489 for host gastrodermis, *p* = 1.000 for algal symbionts).

## Discussion

The association with autotrophic endosymbiotic algae has enabled heterotrophic Cnidaria to thrive in the oligotrophic tropical ocean (Muscatine and Porter 1977; Stanley 2006). The long-term starvation experiment presented here emphasizes the remarkable trophic plasticity that such a symbiosis confers upon these cnidarian holobionts. Because of the highly efficient symbiotic nutrient exchange and recycling, Aiptasia were able to survive without heterotrophic feeding for at least one year. At the same time, starved animals showed clear signs of nutrient limitation, including reduced biomass, host protein content, and symbiont density, underscoring the long-term importance of heterotrophic feeding for body mass maintenance and growth.

### Autotrophic nutrient recycling can sustain the cnidarian-algal symbiosis for extended periods of time

Recent work suggests that the lack of heterotrophic feeding could shift the cnidarian-algal symbiosis towards parasitic interactions that reduce the capacity of the host to survive starvation (Peng et al. 2020). However, here we show that, even after one year of complete heterotrophic starvation, the translocation of photosynthates by algal symbionts remained sufficient to maintain the basal metabolic requirement of the host. Indeed, patterns of host ^13^C enrichment (Fig. 2A-C) were not affected by heterotrophic starvation indicating that photosynthate availability for the host was not impaired despite an 85 % reduction in algal symbiont biomass in the holobiont (Fig. 1B). Combined with the pronounced increase in the density of lipid bodies in the host tissue of starved Aiptasia (Fig. S2), this implies that carbon translocation by individual algal cells must have significantly increased in response to heterotrophic starvation. Indeed, we observed a 50 % increase in ^13^C enrichment and an increased density of lipid bodies among the algal symbionts in starved animals (Fig. 2C), clearly indicating enhanced photosynthetic performance and excess fixed carbon availability required for higher relative translocation rates. Similar, albeit less pronounced, trends were previously reported in a three-month starvation experiment using Aiptasia (Davy and Cook 2001). These authors proposed that the increase in algal photosynthetic performance in starved animals was the result of reduced intra-specific competition for CO_2_. Indeed, reduced algal symbiont densities likely reduce competition for CO_2_ (Rädecker et al. 2017; Krueger et al. 2020). However, in starved animals, this effect could, in part, be masked by the reduced catabolic CO_2_ production in the holobiont due to the lack of heterotrophic prey digestion by the host. Our data point to an additional mechanism that could promote enhanced photosynthate release by algal symbionts in the absence of heterotrophy, namely nitrogen starvation.

### Nitrogen limitation shapes the starvation response of Aiptasia

In the stable symbiosis, low nitrogen availability limits the anabolic incorporation of photosynthates in the algal symbiont metabolism (Rädecker et al. 2021; Cui et al. 2022a, 2022b). This nitrogen limitation is thus not only crucial in regulating algal growth but also ensures the translocation of excess photosynthates to the host (Muscatine and Porter 1977; Falkowski et al. 1984). The host passively modulates *in hospite* nitrogen availability for algal symbionts through ammonium assimilation and production of ammonium in its glutamate metabolism (Rahav et al. 1989; Rädecker et al. 2021; Cui et al. 2022a). Here, we found a proportional decline of algal symbiont density and host protein content in heterotrophically starved Aiptasia (Fig. 1C,D). Given that both algal growth and host protein synthesis depend on nitrogen availability this suggests that starvation caused severe nitrogen limitation, which is consistent with previous work documenting increases in the carbon-to-nitrogen ratio and lipid content of Symbiodiniaceae in unfed Aiptasia (Cook et al. 1988; Cook and Muller-Parker 1992; Muller-Parker et al. 1996). Strongly reduced nitrogen availability could thus drive the enhanced translocation of photosynthates by the algal symbionts observed here and explain the long-term survival of Aiptasia during heterotrophic starvation.

Interestingly, the reduced nitrogen availability did not cause changes to the ammonium assimilation rates by either symbiotic partner; indeed both continued to efficiently assimilate ammonium from the surrounding seawater in the absence of heterotrophic nutrients. Because ammonium assimilation depends on the availability of carbon backbones from the TCA cycle (Cui et al. 2022b), this observation also suggests that starved holobionts did not experience severe carbon limitation. Yet, the starved holobionts showed severe shrinkage and a significant decline in biomass indicative of malnutrition in the present study. Under the current experimental conditions, environmental ammonium assimilation was thus not sufficient to fulfill the nitrogen requirements of the holobiont. The availability of seawater ammonium was possibly limited by the rate of water exchange in our experiment (once per week as for the entire animal culture stock). It is thus plausible that higher ammonium concentrations would have allowed the heterotrophically starved Aiptasia to maintain a larger fraction of their original biomass. Yet, in the environmental context of the oligotrophic ocean, photosymbiotic animals are likely similarly limited in their access to environmental ammonium (O’Neil and Capone 2008). In this context, our findings illustrate the importance of heterotrophic feeding by the host for the long-term maintenance of the cnidarian-algal symbiosis biomass. While symbiotic nutrient exchange and recycling may be sufficient to cover the carbon and energy demands of the symbiotic partners on a time scale of at least one year, heterotrophic feeding is not only required for long-term survival but also required for propagation and net growth of the holobiont.

## Conclusion

This study has illustrated the substantial trophic plasticity of the cnidarian-algal symbiosis: Aiptasia survived for an entire year in the complete absence of heterotrophic feeding. Our findings reveal that efficient symbiotic nutrient exchange and recycling are sufficient to sustain the basic metabolic requirements of both symbiotic partners over extended periods of time. Yet, under long-term exposure to highly oligotrophic conditions, the assimilation of environmental inorganic nitrogen is not sufficient to support the nutritional requirements of the holobiont, and heterotrophic feeding represents an essential source of nitrogen for holobiont growth.,Mixotrophy thereby provides a nutritional advantage to photosymbiotic cnidarians that in part explains their ability to outcompete other organisms restricted to either autotrophic or heterotrophic nutrient acquisition under oligotrophic conditions.

## Acknowledgments

We are grateful to Dr. Claudia Pogoreutz and Gaëlle Toullec for their help with animal culture maintenance. We thank Jean Daraspe and Dr. Cristina Martin-Olmos for their help with sample processing and SEM imaging as well as Dr. Stéphane Escrig and Florent Plane for their assistance and support with NanoSIMS measurements. We thank the editor/recommender as well as the two reviewers for their constructive feedback that significantly improved the manuscript.

## Supplementary material for

**Fig. S1.**
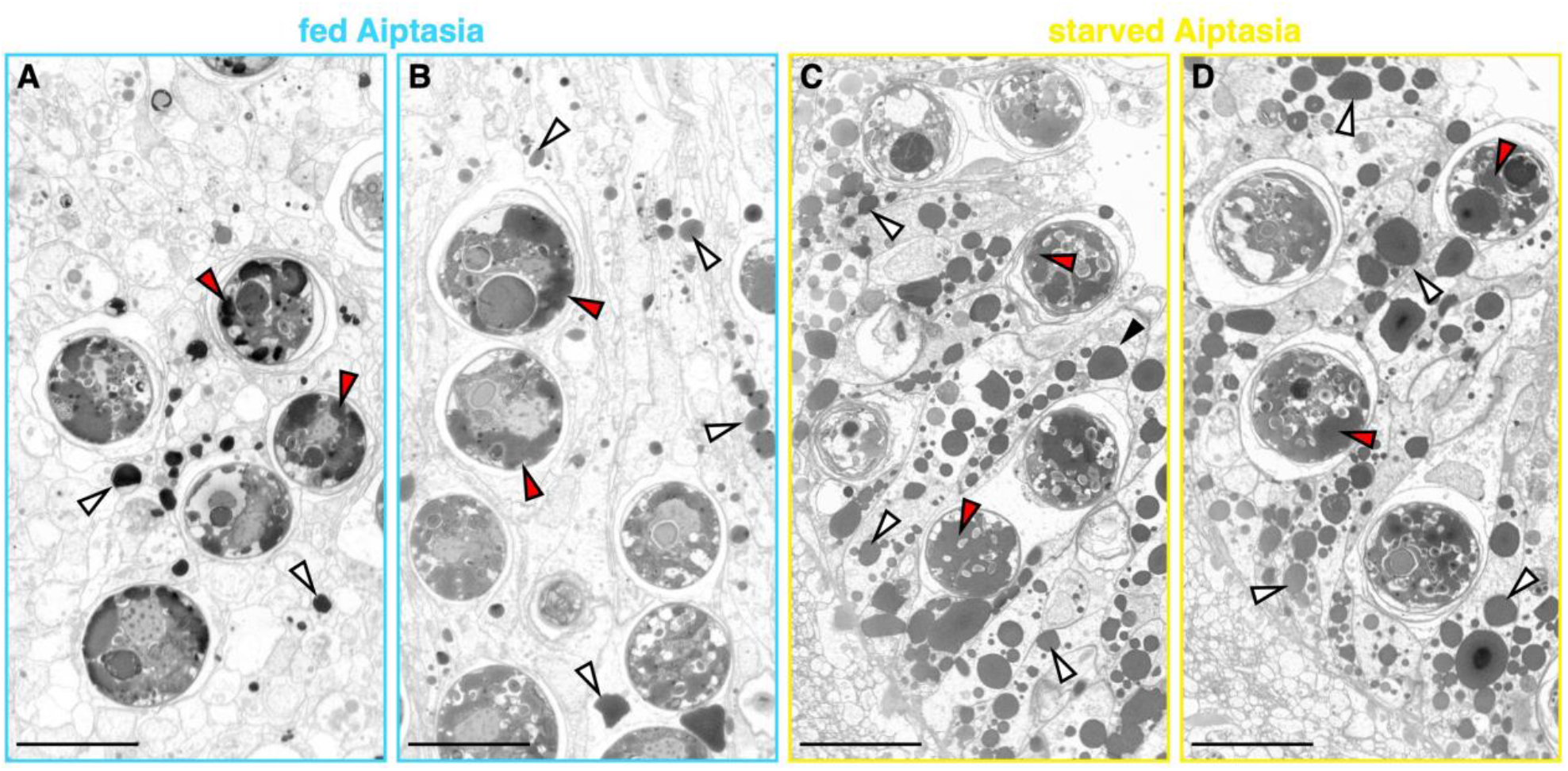
SEM images of semi-thin sections of gastrodermal regions of tentacles from (A,B) regularly fed and (C,D) starved Aiptasia. White triangles point to lipid bodies in the host gastrodermis; red triangles point to lipid bodies in the algal symbionts. Scale bars are 10 μm.

**Fig. S2.**
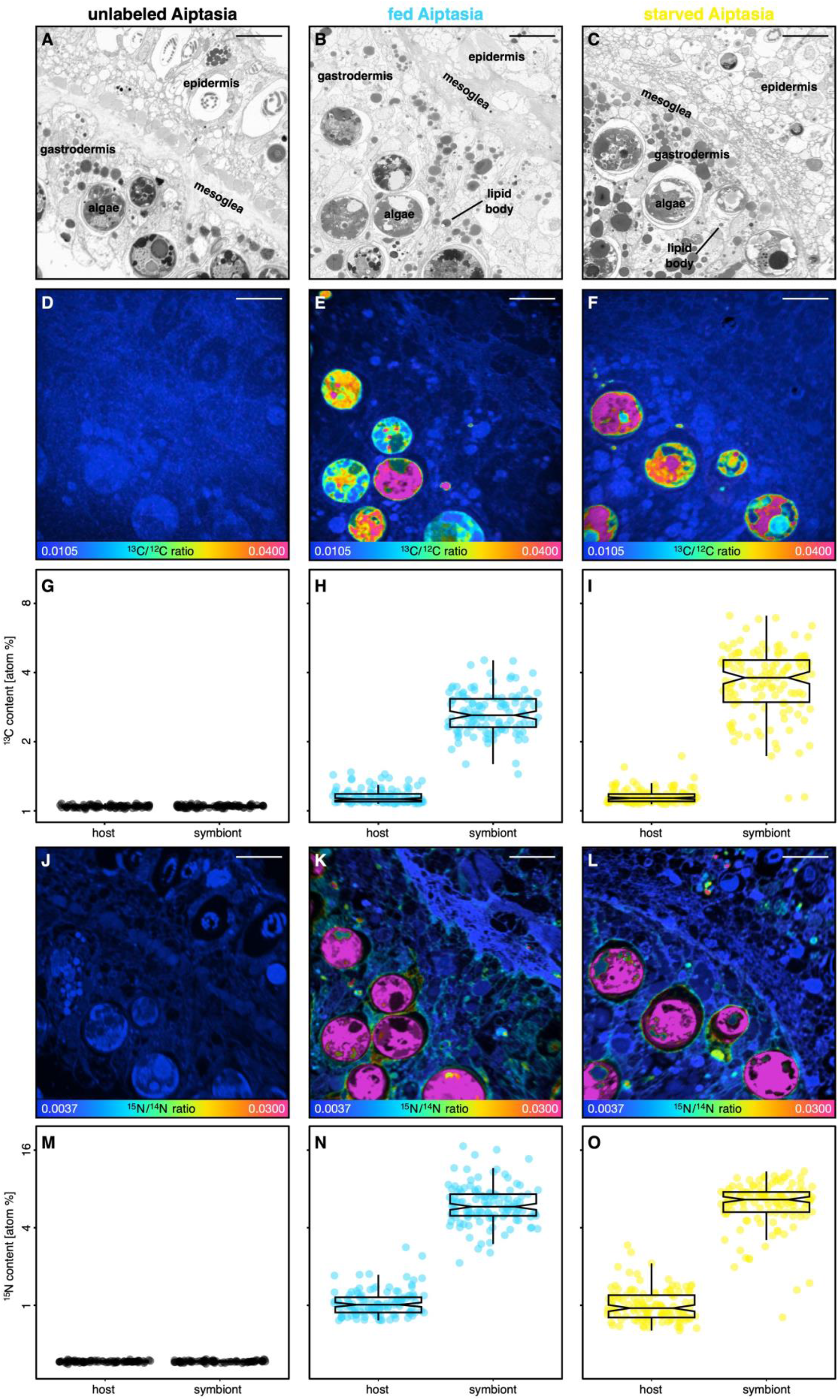
Correlative SEM and NanoSIMS imaging of symbiotic carbon and nitrogen cycling in unlabeled, fed, and starved Aiptasia. **(A-C)** Representative SEM images of tentacle semi-thin sections of unlabeled, regularly fed and starved Aiptasia, respectively. **(D-F)** Corresponding NanoSIMS images illustrating H^13^CO_3_- assimilation and translocation as ^13^C/^12^C isotope ratio maps. **(G-I)** Boxplots and data points of ^13^C content for the host gastrodermis and the algal symbiont cells. **(J-L)** Corresponding NanoSIMS images illustrating ^15^NH_4_^+^ assimilation as ^15^N/^14^N isotope ratio maps. **(M-O)** Boxplots of ^15^N enrichment for the host gastrodermis and the algal symbiont cells. NanoSIMS ratio maps are shown as hue saturation images with blue representing no/low enrichment and pink representing the highest level of enrichment. Note the logarithmic scale for **G-I** and **M-O**. Scale bars are 10 μm. For the unlabeled Aiptasia, 80 regions of interest across two Aiptasia replicates were analyzed. For the other treatments, 120 regions of interest across three Aiptasia replicates were analyzed per symbiotic partner.

